# Spontaneous Histone Exchange Between Nucleosomes

**DOI:** 10.1101/2023.05.09.540004

**Authors:** Subhra Kanti Das, Mai Thao Huynh, Tae-Hee Lee

## Abstract

The nucleosome is the fundamental gene-packing unit in eukaryotes. Nucleosomes comprise ∼147 bp DNA wrapped around an octameric histone protein core composed of two H2A-H2B dimers and one (H3-H4)_2_ tetramer. The strong yet flexible DNA-histone interactions are a physical basis of the dynamic regulation of genes packaged in chromatin. The dynamic nature of DNA-histone interactions implies that nucleosomes dissociate DNA-histone contacts transiently and repeatedly. This kinetic instability may lead to spontaneous nucleosome disassembly or histone exchange between nucleosomes. At a high nucleosome concentration, nucleosome-nucleosome collisions and subsequent histone exchange would be a more likely pathway, where nucleosomes act as their own histone chaperone. The spontaneous histone exchange would serve as a mechanism for maintaining the overall chromatin stability although it has never been reported. We employed three-color single-molecule FRET (smFRET) to demonstrate that histone H2A-H2B dimers are exchanged spontaneously between nucleosomes and that the time scale is on a few tens of seconds at a physiological nucleosome concentration. The rate of histone exchange increases at a higher monovalent salt concentration, with histone acetylated nucleosomes, and in the presence of histone chaperone Nap1, while it remains unchanged at a higher temperature, and decreases upon DNA methylation. These results support histone exchange via transient and repetitive partial disassembly of the nucleosome and corroborate spontaneous histone diffusion in a compact chromatin context, modulating the local concentrations of histone modifications and variants.

## INTRODUCTION

The nucleosome is the basic gene-packing unit in eukaryotes. Nucleosomes further compact into chromatin, and eventually into a chromosome. A nucleosome core particle comprises a ∼147 base-pair (bp) DNA fragment wrapped around a histone protein core in 1.67 left-handed superhelical turns^1,2^. The histone protein core is composed of two H2A-H2B dimers and one (H3-H4)_2_ tetramer, forming the characteristic octameric structure^1^. The negatively charged DNA backbone interacts strongly with the positively charged histone surface to compact genes, casting high energy barriers for DNA access^2^. On the other hand, the nucleosome structure is flexible to allow for dynamically regulated DNA access. For instance, the DNA termini of the nucleosome undergo ‘breathing’ motions, rendering transient and repetitive DNA accessibility^3–5^.

These strong yet flexible DNA-histone interactions are necessary to enable dynamic gene regulation while maintaining the structural integrity of chromatin. However, the dynamic nature of the nucleosome structure leads to its disassembly at a sub-nanomolar concentration^3,6^. The kinetic instability suggests that nucleosomes undergo constant, repetitive, and transient partial disassembly^6^. A cryo-EM structural study showed that many nucleosomes have ∼15 bp of DNA unwrapped at the entry/exit region and that this partially disassembled state can be stabilized by internal rearrangement of histone H2A-H2B dimers^7^. At a high concentration of nucleosomes, such transiently disassembled nucleosomes may interact with each other to form inter-nucleosomal DNA-histone contacts. To support this hypothesis, a recent study reported di-nucleosome stacking between two partially disassembled nucleosomes through inter-nucleosomal DNA-histone interactions^8,9^. Under this hypothesis, nucleosomes would function as their own histone chaperone, stabilizing the entire population of nucleosomes at a high nucleosome concentration as in chromatin. Histone chaperones mediate DNA-histone interactions to help assemble or disassemble nucleosomes depending on the context and eventually drive a DNA-histone mixture to their thermodynamic equilibrium^10^. Histone chaperones typically contain an acidic surface to compete against DNA for histone binding^11–13^. In this regard, naked DNA may also act as histone chaperone^14^. The histone chaperone function of the nucleosome would result in stabilized nucleosomes at the cost of freely and spontaneously exchanged histones, although such spontaneous histone exchange has never been reported. To support this notion, exchange of nucleosomal H2A-H2B with exogenous histone H2A-H2B has been reported^15,16^. Histone exchange among nucleosomes on a relevant timescale would imply that histone modifications and variants can diffuse and redistribute themselves spontaneously in chromatin after serving their purpose.

Some nucleosome modifications mediate gene regulation by altering the DNA-histone dynamics in the nucleosome^17–19^. DNA methylation at CpG dinucleotide sequences has been reported to induce attenuated DNA-histone dynamics^20–22^. Various post-translational modifications (PTM) of histones such as phosphorylation, acetylation, and methylation have been coupled to gene regulation^23–27^. In particular, histone H3 acetylation at K56 (H3K56ac) has been associated with elevated DNA accessibility by facilitating DNA-histone motions in the nucleosome^17,22,28–30^.

Here we investigated spontaneous histone H2A-H2B dimer exchange between nucleosomes under various conditions with three-color single-molecule FRET (smFRET). Our results indicate that histone H2A-H2B dimers are exchanged spontaneously between nucleosomes at a rate corresponding to a few tens of seconds exchange timescale at physiological nucleosome and monovalent salt concentrations. The exchange reaction is facilitated at a higher monovalent salt concentration or with H3K56ac^6,16^, supporting spontaneous, transient, and repetitive partial disassembly of the nucleosome as the underlying mechanism. We investigated the effect of histone chaperone Nap1 to support that it facilitates histone exchange, thereby catalyzing histone equilibrium among nucleosomes in chromatin. We also found that CpG methylation significantly lowers the rate of histone exchange, suggesting its role in suppressing DNA accessibility and maintaining chromatin integrity. The rate constants remain unchanged at 25 °C from those at 4 °C, indicating that the DNA-histone interactions are not disturbed further enough to make a difference under these two conditions. These results support a model where nucleosomes are stabilized in chromatin by chaperoning themselves, and histone modifications and variants in H2A-H2B are spontaneously redistributed and equilibrated on a timescale of a few tens of seconds in chromatin.

## EXPERIMENTAL PROCEDURES

### Construction of nucleosomal DNA

Single-stranded DNA (ssDNA) oligonucleotides were purchased from Integrated DNA Technologies (IDT Inc., Coralville, IA). The sequences are listed in Table S1. Nucleosomal DNA was constructed by following previously published protocols^30,31^. Briefly, each nucleosomal DNA construct was prepared by annealing and ligating the oligonucleotides. Oligonucleotides F2 and F3 (Table S1) were labeled with Cy5 and Cy3 fluorophores respectively, via conjugation between a C6-amino linker and an NHS-ester functionalized fluorophore.

### Histone H3K56 acetylation and histone chaperone Nap1 preparation

Histone H3 K56C/C110A double mutant (*Xenopus laevis*) was purchased from The Histone Source (Colorado State University). The histone was dissolved in 200 μL of the reaction buffer containing 0.2 M sodium acetate (pH 4), 6 M guanidine–HCl, 7 mM L-glutathione, 50 mM N-vinyl acetamide, 100 mM dimethyl sulfide, and 5 mM VA-044 (2,2-[azobis(dimethyl methlene)]bis(2-imidazoline)-dihydrochloride) to the final concentration of 1 mM as was previously published^32^. The reaction was carried out in complete darkness for 2 hours at 70 °C in a water bath. The resulting histone H3 has a thioether-linked sidechain of acetylated lysine at residue C56, often denoted by H3K_C_56ac. We will denote this acetylation by H3K56ac for simplicity. The product was dialyzed against deionized water three times for 2 hours, 4 hours, and overnight, and then lyophilized. The solid form of the acetylated histone protein was stored at -80 °C. 6xHis-Yeast nucleosome assembly protein 1 (Nap1) was expressed in *E. coli* BL21(DE3) pLysS and purified with Ni-NTA beads (Thermo Fisher Scientific, Waltham, MA) as was reported in a previous publication^33^.

### Preparation of CpG methylated DNA

CpG methylation on the DNA was carried out by following standard protocols for the CpG methyltransferase enzyme M.SssI (New England Labs Inc.). Briefly, fluorophore labeled or unlabeled nucleosomal DNA was mixed with S-adenosylmethionine (SAM) in the methyltransferase reaction buffer (50 mM NaCl, 10 mM Tris-HCl (pH 7.9), 10 mM MgCl_2,_ 1mM DTT) and the M.SssI enzyme. The reaction mixture was kept at 37 °C for 4 hours. The reaction was stopped by heating the mixture at 65 °C for 20 minutes.

### Preparation of histone octamer(s), histone labeling, and nucleosome reconstitution

The unmodified core histone octamer (*Xenopus laevis*) was purchased from The Histone Source (Colorado State University). The stock was diluted to the final concentration of 2 mM for nucleosome reconstitution in a buffer (10 mM Tris-HCl (pH 7.5), 2M NaCl, 5 mM 2-mercaptoethanol, 1 mM EDTA). For labeled histone core, histone proteins H2A, H2B T112C, H3, and H4 (*Xenopus laevis*) were purchased from The Histone Source (Colorado State University). Histone H2B T112C was labeled by published protocols^30^. Briefly, H2B T112C mutant was unfolded in an unfolding buffer (20 mM Tris-HCl (pH 7.5), 7 M guanidium hydrochloride, 10 molar excess TCEP) followed by overnight incubation at 4 °C with 20 molar excess sulfo-maleimide Cy5.5 dye (Lumiprobe, Baltimore, MD). The excess dye was removed through several PD-10 desalting columns (GE Healthcare). The labeling efficiency was 53.7% according to a UV-Vis measurement. All histone proteins were unfolded in an unfolding buffer (20 mM Tris-HCl (pH 7.5), 7 M Guan-HCl (guanidium hydrochloride), 1 mM DTT) for at least 30 minutes. Histone H2A-H2B and (H3-H4)_2_ were then assembled separately by dialyzing against refolding buffer (2M NaCl, 10 mM Tris-HCl (pH 7.5), 1mM EDTA, and 5mM BME) for 2 hours, 4 hours, and overnight. The octamer was purified with a size exclusion column (HiLoad 16/600 Superdex 200pg, GE Healthcare). The presence and purity of H2A-H2B and (H3-H4)_2_ were confirmed by SDS-PAGE.

Nucleosome reconstitution was carried out by mixing stoichiometric amounts of DNA and histones according to the previously reported protocols^34^. We used salt gradient dialysis method in a dialysis cup (Slide-A-Lyzer MINI Dialysis Device, 7K MWCO, Thermo Fisher Scientific) using 1× TE (pH 8.0) buffer with stepwise decreasing salt concentrations of 1200, 850, 600, 400, 200, and 10 mM NaCl. The purity of all nucleosome sets was confirmed with native-PAGE (Fig. S1).

### Microscope slide preparation

Pre-drilled microscope quartz slides were purchased from G. Finkenbeiner Inc. (Waltham, MA). The cleaned slides were constructed with five flow channels and passivated with a lipid bilayer following published protocols^30,35^. Briefly, a sub-monolayer of biotin-PEG-silane was coated (MW 3400, Laysan Bio, Arab, AL) on a carefully and thoroughly cleaned quartz microscope slide followed by deposition of a lipid bilayer. For preparation of the lipid solution, dried lipid vesicles made of 1,2-Dioleoyl-sn-Glycero-3-phosphoethanolamine-N-[Methoxy(Polyethyleneglycol)-5000] (ammonium salt) (DOPC) (Avanti Polar Lipids, product number 880230, Alabaster, AL) were suspended in a buffer containing 10 mM Tris-HCl (pH 7.8) and 100 mM NaCl. The suspension was subjected to 1 min sonication/cooling cycles on ice with a tip-based (Branson

Ultrasonics, part no #101-148-062) ultrasonic cell disruptor (Sonifier 550, Branson Ultrasonics) until the lipid suspension became transparent. Once the suspension became transparent, the lipid vesicles were extruded through a 100 nm pore-sized polycarbonate membrane filter (Avanti Polar Lipids, Alabaster, AL). Flow channels on the surface of a biotin-PEG coated microchannel were injected with 1 mg/mL concentrated lipid solution and incubated for 45 minutes to deposit a bilayer (Fig. 1C). Following the lipid bilayer deposit, 40 μL of 0.1 mg/mL streptavidin solution was injected into the flow channels and incubated for 15 mins for biotin-streptavidin conjugation. This conjugation is to immobilize the nucleosome samples on the slide surface for subsequent smFRET measurements.

**Figure 1.**
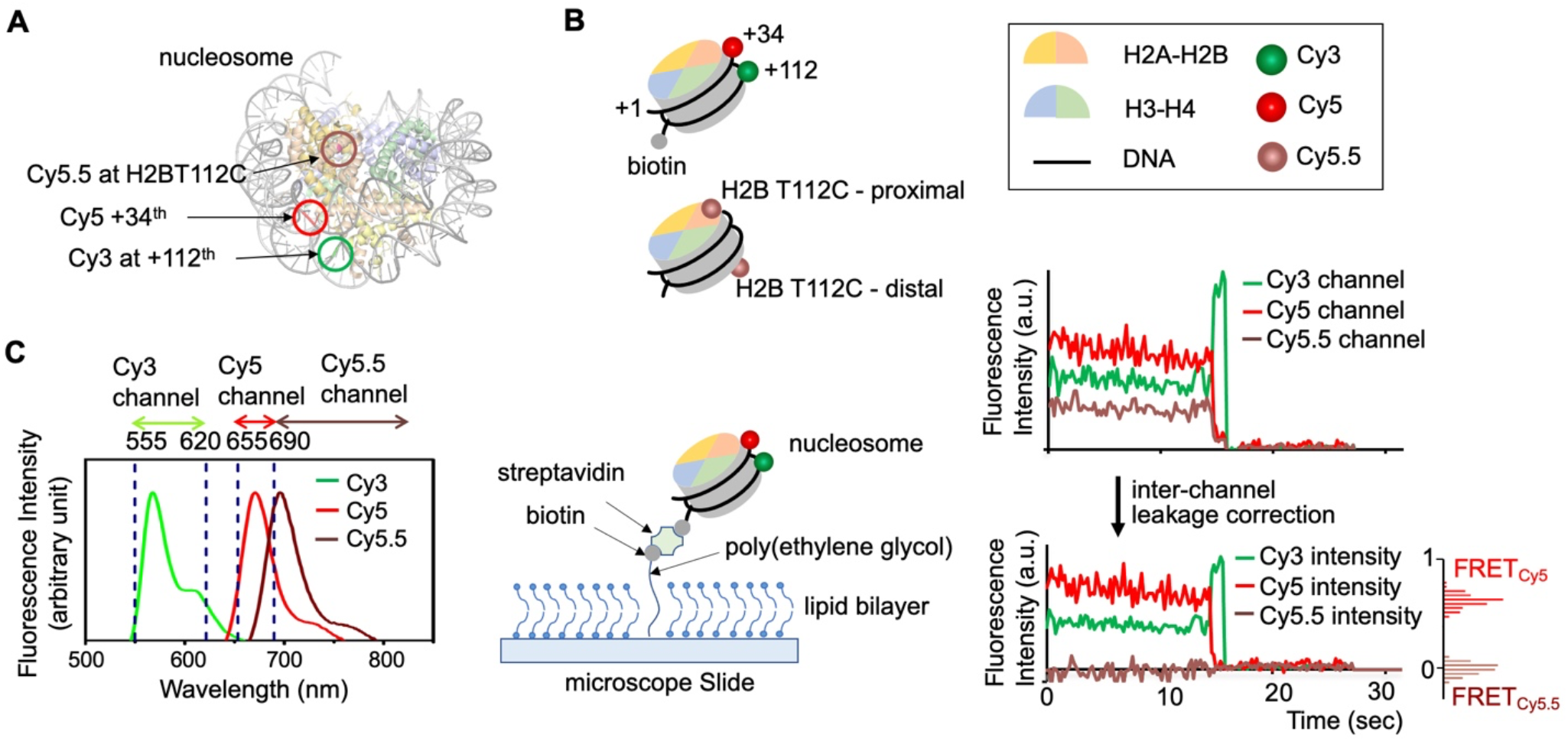
Three-color single-molecule FRET setup to monitor histone H2A-H2B exchange between nucleosomes. (A) A nucleosome structure (PDB ID: 3LZ0) is shown with three fluorophore labeling positions marked. (B) Fluorophore labeling schemes are illustrated for two types of nucleosomes, where one is labeled with Cy3 and Cy5 at the +112^th^ and +34^th^ nucleotides counted from the nucleosome entry, and the other is labeled with Cy5.5 at histone H2B T112C. (C) Separations of the three spectral regions for fluorescence detection (left), microscopic measurement setup (center), and representative fluorescence intensity traces for an intact nucleosome labeled with Cy3 and Cy5 before and after inter-channel leakage correction (right) are shown. Fluorescence intensities are in arbitrary unit (a.u.). The inter-channel leakage corrected traces confirm high FRET_Cy5_ and zero FRET_Cy5.5_ values. Their histograms are shown on the side.

### Sample preparation and smFRET measurements for dimer-exchange reaction

Dimer exchange reaction(s) were carried out at 4 °C after mixing DNA labeled-nucleosomes with histone-labeled nucleosomes at a 1:1 ratio. The final concentration of the nucleosomes in the stock was 200 nM each, resulting in a total concentration of 400 nM. These are the *reaction stock(s)* for the dimer-exchange reactions. Upon preparing a reaction stock, small aliquots were taken out and each aliquot was incubated for 0, 1, 2, 4, 6, 8, 12, 18, or 24 hours before measuring the fraction of histone exchanged nucleosomes. For the measurement, each aliquot was diluted to a concentration of ∼ 70 – 100 pM in an imaging buffer containing 10 mM Tris-HCl (pH 8.0), 1 mM Trolox, 0.1 mg/ml of BSA, 2 mM protocatechuic acid (PCA) and 0.2 U/ml protocatechuate-3,4-dioxygenase (PCD). A total of 4 – 10 independent measurements were made per time point. For the low salt reaction condition, 10 mM NaCl was used, whereas for the high salt condition, 50 mM NaCl and 150 mM KCl were used. Room temperature exchange reactions were carried out at 25 °C at 10 mM NaCl. Nap1-mediated exchange reactions were carried out at 400 nM Nap1 (nucleosome: Nap1 = 1:1) at 10 mM NaCl.

### Three-color smFRET setup

Three-color smFRET measurements were taken with a custom-built EMCCD-based TIRF setup as was reported previously^36^. Briefly, a 532 nm green laser (CrystaLaser, Reno, NV) and a 635 nm red laser (CrystaLaser, Reno, NV) were used for FRET donor and acceptor excitation respectively. The surface of a flow channel was illuminated with a green laser in a prism-coupled TIR geometry and the fluorescence images were recorded with an EMCCD camera (IXON Ultra 897, Oxford Instruments), followed by a brief dark period and excitation with 635 nm red laser to verify the existence of Cy5 and/or Cy5.5. The signal integration time for fluorescence imaging was 200 ms. In this one-donor (Cy3) two-acceptor model (Cy5 and Cy5.5), the fluorescence signals from the nucleosomes were spectrally separated into three signals (Cy3, Cy5, and Cy5.5 channels) (Fig. 1C). The relative intensities from a Cy3-Cy5 pair in an intact nucleosome in the three spectral regions are shown in figure 1C. The three-color scheme ensures that we count only properly assembled nucleosomes before and after H2A-H2B exchange and that the measurements are free from any errors due to fluorescent contaminants and impurities originated from sample preparation.

### Background correction and inter-channel leakage correction for single-molecule fluorescence intensities

Time traces of the fluorescence intensities in the three spectral channels (Cy3, Cy5, and Cy5.5 channels) were obtained from the single-molecule experiments (Figs. 1 and 2). These raw intensities are compounded by a constant background and inter-channel leakages mainly between the two acceptors Cy5 and Cy5.5 channels. To correct the constant background, we obtained the background intensities in each channel after the fluorophores were photobleached. These constant background values were subtracted from the intensities of the fluorophore.

**Figure 2.**
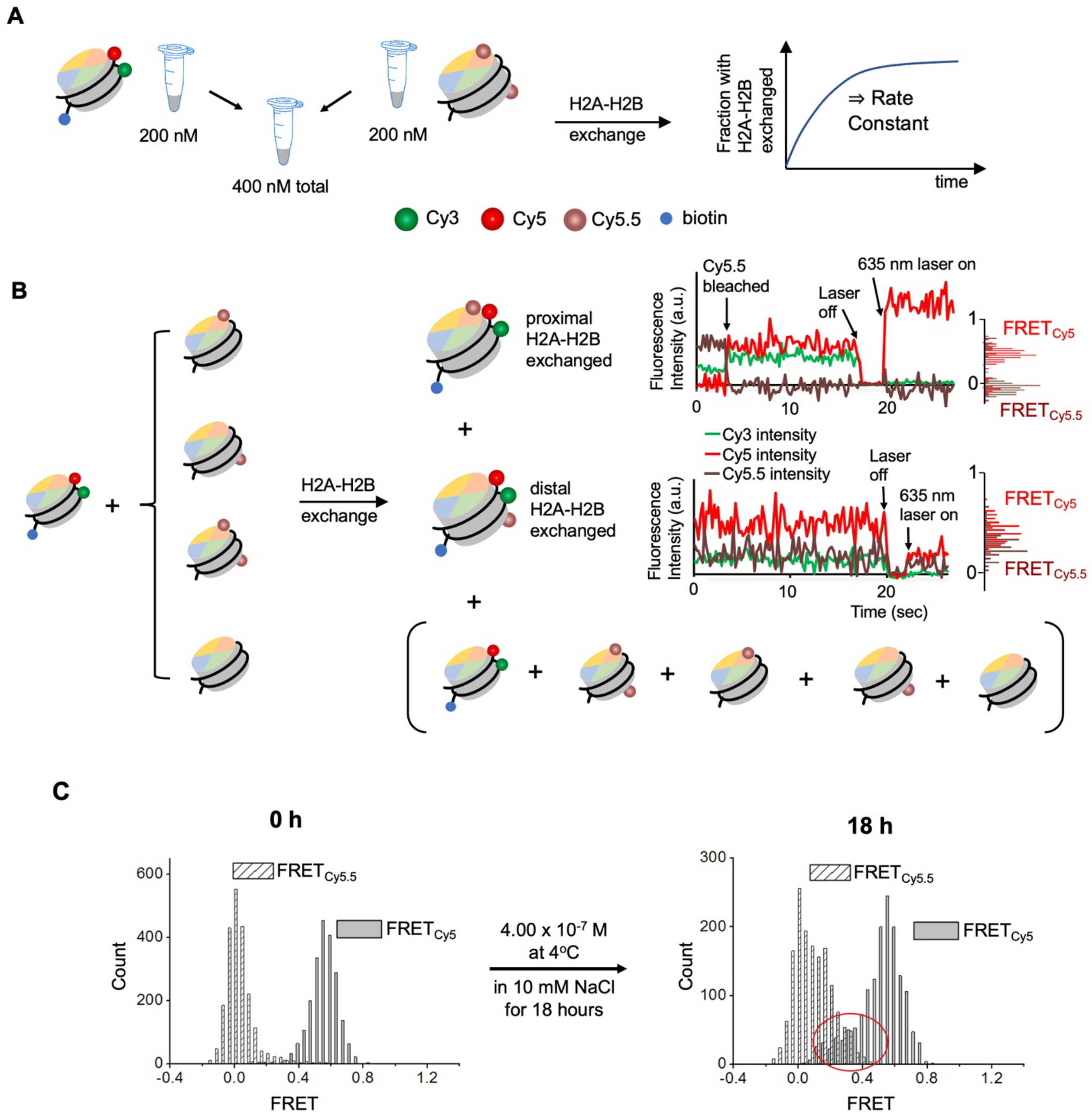
Experimental scheme to monitor the kinetics of histone H2A-H2B exchange between nucleosomes. (A) Two types of nucleosomes differently labeled are mixed at the total concentration of 400 nM under various conditions to monitor the fraction of nucleosomes with H2A-H2B exchanged. (B) The scenarios leading to H2A-H2B exchange detectable with the three-color smFRET setup shown. The reaction will result in nucleosome-entry proximal or distal H2A-H2B exchanged to display observable changes in the FRET signals as shown in the fluorescence intensity traces that are corrected for inter-channel leakage. The fluorescence intensities are in arbitrary unit (a.u.). The histograms of FRET_Cy5_ and FRET_Cy5.5_ are also shown on the side of the traces. In case of entry proximal H2A-H2B exchange, a high FRET_Cy5.5_ and a low FRET_Cy5_ signals are detected, which leads to fast photobleaching of Cy5.5. At the end of an observation time window, the laser is turned off and subsequently, a 635 nm laser is turned on to ensure that the intensity change is due to Cy5.5 photobleaching. In case of distal H2A-H2B exchange, low-mid FRET_Cy5,_ and non-zero FRET_Cy5.5_ signals are observed. The other nucleosomes shown in the parentheses do not generate any Cy5.5 signals. (C) Histograms of FRET_Cy5_ and FRET_Cy5.5_ from the nucleosomes showing zero intensities after photobleaching (single nucleosomes for accurate background correction), longer photobleaching lifetimes (> 10 sec), and a decent signal-to-noise ratio (> 4) at 0 (intact nucleosomes labeled with Cy3 and Cy5) and the 18^th^ hour time points (nucleosomes with Cy3-Cy5 and the entry-distal H2A-H2B exchanged) (8^th^ hour time point for H3 K56 acetylated nucleosomes) time points after mixing. The histograms show the growth of non-zero FRET_Cy5.5_ and lower FRET_Cy5_ counts as marked in the red circled area, confirming histone exchange. Note that the histogram counts cannot be used as a measure of the percent histone exchange as Cy5.5 photobleaches much faster than Cy5.

The fluorescence from an acceptor (Cy5 or Cy5.5) is leaked into the other acceptor channel. To compute their intensities without the contribution from the other acceptor, we first obtained two leakage factors 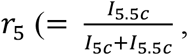 where *I*_5C_ and *I*_5.5C_ are the intensities of Cy5 in the Cy5 and Cy5.5 channels, respectively) and 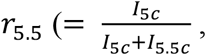 where *I*_5C_ and *I*_5.5C_ are the intensities of Cy5.5 in the Cy5 and Cy5.5 channels, respectively). The value of *r*_5_ represents the fraction of Cy5 emission registered in the Cy5.5 channel and can be obtained by measuring the intensities of Cy5 in the Cy5 and Cy5.5 channels. The value of *r*_5.5_ represents the fraction of Cy5.5 emission registered in the Cy5 channel and can be obtained by measuring the intensities of Cy5.5 in the Cy5 and Cy5.5 channels. We made multiple measurements of the *r*_5_ and *r*_5.5_ values and used the average values of 0.31 ± 0.02 and 0.27 ± 0.04 for *r*_5_ and *r*_5.5_, respectively for leakage correction. Let the true fluorescence intensities of the Cy5 and Cy5.5 without leakage be *I*_5_ *and I*_5.5_. Writing *I*_5_ *and I*_5.5_ in terms of the apparent intensities from the 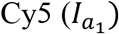 and Cy5.5 channels 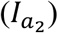,

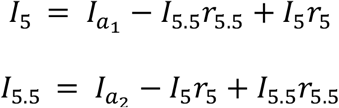

Solving these two equations for the true intensities *I*_5_ and *I*_5.5_ results in

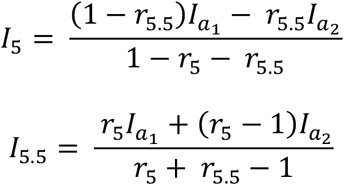

These leakage-corrected intensities were used to construct the time traces of Cy5 and Cy5.5 intensities and their FRET efficiency histograms in figure 2 and figure S3 where the FRET efficiencies for Cy5 (*FRET*_Cy5_) and Cy5.5 (*FRET*_Cy5.5_) are defined as

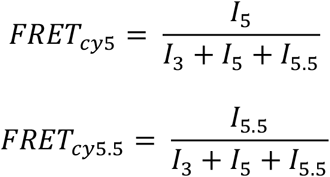

where *I*_3_, *I*_5_, and *I*_5.5_ are the intensities of Cy3, Cy5, and Cy5.5 after background and leakage correction.

### Identifying dimer-exchanged nucleosomes based on the fluorescence intensity signatures

To identify a dimer-exchanged nucleosome, Cy3/Cy5/Cy5.5 relative intensity levels were examined from the surface immobilized nucleosomes (Fig. 2). In the entry-proximal H2A-H2B exchanged case, the distances between the three fluorophores result in near 100 % FRET funneling to Cy5.5 (Fig. S2A). We modeled this scenario with two 15-bp DNA fragments labeled with Cy3, Cy5, and Cy5.5 with similar inter-dye distances (Fig. S2A). The three fluorophore intensities are shown in figure S1C that are similar to what we observed from some of the histone-exchanged nucleosomes, confirming proximal H2A-H2B exchange (Fig. 2B). In the entry-distal H2A-H2B exchanged case, the distances between the fluorophores should result in strong FRET to both Cy5 and Cy5.5 from Cy3 (Figs. S2B and S2C). We used another pair of 15-bp DNA fragments labeled with the same fluorophores to simulate the scenario (Fig. S2B). The relative intensities of the three fluorophores are very similar to some of the histone-exchanged nucleosomes, confirming distal H2A-H2B exchange (Figs. 2B and S2C).

### Extracting the rate constant of histone exchange

We counted the numbers of intact nucleosomes and distal H2A-H2B exchanged nucleosomes as described above and calculated the fractions of histone exchange at various time points. The time course of the fractions was fitted with an equation approximating the kinetics to extract the histone exchange rate constant. Considering partial labeling of H2A-H2B and near complete labeling of DNA, figure S3A illustrates nucleosome species with all possible fluorophore combinations. Figure S3B lists histone exchange between all possible such labeled pairs of the nucleosomes with the same rate constant *k*. These reactions lead to the differential rate equation for the production of trackable H2A-H2B exchanged nucleosomes (i.e. species B in figure S3A) as follows:

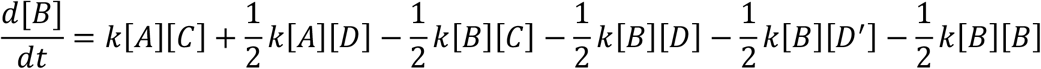

Let [B] be its mole fraction *x*, then the mole fractions of the other species are as follows in the beginning.

[A] = 0.5 – *x*, [B’] ≈ 0, [b] = [e] = *x*/2, [C] = (0.5 × *F* ^2^ – *x*), [D]= *x* + eff(1-*F*), [d] = [f] = [*x* + F(1-*F*)]/2, and [D’] = 0.5(1 - *F*)^2^, where *F* is the H2A-H2B labeling efficiency that is 53.7 % according to our UV-Vis measurement. Writing the differential rate equation in terms of mole fractions is as follows:

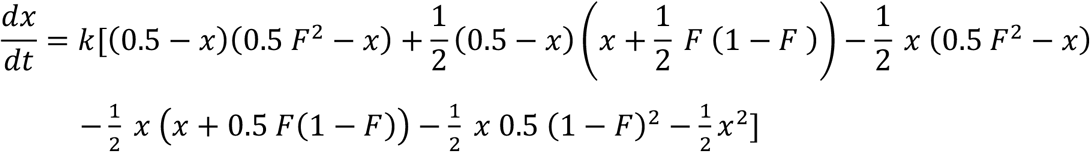

Note that, we approximate the mole fraction of both H2A-H2B exchanged as negligible (i.e. [B’] ≈ 0), which should be reasonable during an early time period.

We counted the number of species e, the distal H2A-H2B exchanged nucleosomes. Solving the differential equation for *x(t)*/2 (= mole fraction of e as a function of time) leads to the following equation.

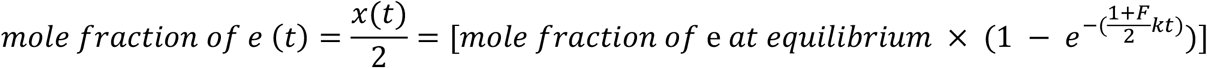

The mole fraction of species e at equilibrium is the probability of having the entry-proximal H2A-H2B unlabeled and the other H2A-H2B labeled in a nucleosome (= (1-0.537/2) × 0.537/2 = 0.196, where 0.537/2 is the mole fraction of labeled H2A-H2B in the reaction mix and 0.537 is the labeling efficiency of H2A-H2B). Therefore, the approximate mole fraction [e] as a function of time is as follows.

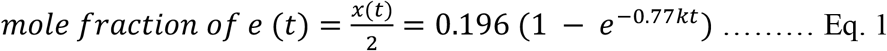

## RESULTS

### Histone H2A-H2B dimers are spontaneously exchanged between nucleosomes

Monitoring H2A-H2B exchange between nucleosomes as a function of time is challenging with an ensemble averaging method due to various sources of heterogeneity such as contaminants and impurities from sample preparation and labeling. We constructed two types of differently labeled nucleosomes to investigate the rate of histone H2A-H2B exchange between nucleosomes (Fig. 1). One type of the nucleosomes is labeled with Cy3 and Cy5 at DNA as a FRET pair (Fig. 1B). This nucleosome is also labeled with biotin for surface immobilization. The fluorescence signals from Cy3 and Cy5 were divided into three spectral channels and imaged on a camera as is shown in figure 1C. The FRET histograms in figure 1B constructed with the fluorescence intensities recorded in the three spectral channels confirm ∼0.6 FRET_Cy5_ and zero FRET_Cy5.5_. The other type of the nucleosomes has H2A-H2B labeled with Cy5.5 and no labels on the DNA (Fig. 1B). After equal amounts of both nucleosomes were mixed to a final concentration of 200 nM each at 10 mM NaCl, the fraction of nucleosomes with H2A-H2B exchanged at 4 °C was monitored based on the smFRET signals (Fig. 2). Upon H2A-H2B exchange between the two types of the nucleosomes, a FRET_Cy5.5_ signal arises. The Cy5.5 fluorophore labeled at the H2A-H2B proximal to the nucleosome entry would result in a high FRET_Cy5.5_ and strong fluorescence from Cy5.5, which leads to its fast premature photobleaching before proper observation (Fig. 2B). We constructed a mimetic 3-color smFRET system with DNA (Fig. S2B) to confirm the relative intensity levels of Cy3, Cy5, and Cy5.5 in case of proximal H2A-H2B exchange. The Cy5.5 fluorophore labeled at the H2A-H2B distal to the nucleosome entry will result in a low Cy3 signal and a mid-high Cy5 and Cy5.5 signals (Fig. 2B), as is confirmed with another DNA mimetic (Fig. S2C). The histograms of FRET_Cy5_ and FRET_Cy5.5_ in the case of distal H2A-H2B dimer exchange (Fig. 2B) show non-zero FRET_Cy5.5_ counts and lower FRET_Cy5_ values than those from the intact nucleosome before exchange (Fig. 1B). We counted the distal dimer exchanged nucleosomes since counting the proximal dimer exchanged nucleosomes results in a large error due to their premature photobleaching. Histograms of FRET_Cy5_ and FRET_Cy5.5_ from the nucleosomes showing zero intensities after photobleaching (single nucleosomes for accurate background correction), longer photobleaching lifetimes (> 10 sec), and a decent signal-to-noise ratio (> 4) before and after distal H2A-H2B exchange are shown in figure 2C and figure S4. Note that the histograms cannot be used as a measure of the efficiency of histone exchange as Cy5.5 photobleaches much faster than Cy5. According to our experience, it is typical that fluorophores labeled on protein photobleach much faster than those on nucleic acids. Instead, we counted the number of nucleosomes displaying the sign of distal H2A-H2B exchange (i.e. non-zero FRET_Cy5.5_ with concomitantly lowered FRET_Cy5_) at various time points during the reaction and computed the fractions of H2A-H2B exchanged nucleosomes. The fractions are plotted against time and fitted with Eq. 1 to obtain the exchange rate constant (Fig. 3A). The rate constant at 10 mM NaCl at 4 °C is 356 ± 64 M^-1^s^-1^. This rate corresponds to 40 ± 7 sec H2A-H2B exchange time at a 70 μM nucleosome concentration as in a human nucleus (3.0 × 10^6^ nucleosomes per 690 μm^3^ nucleus volume)^37^.

**Figure 3.**
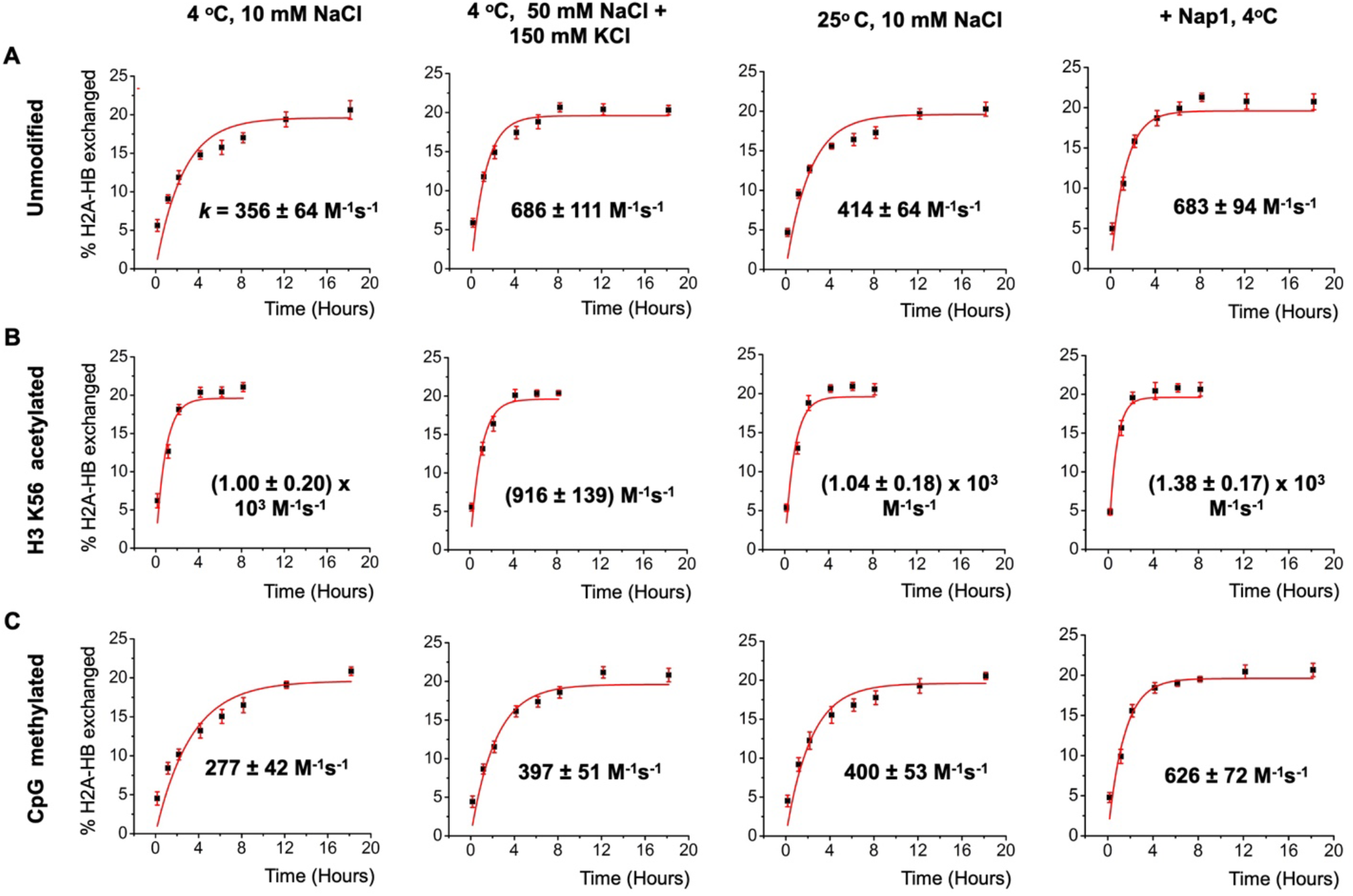
Kinetics of spontaneous histone H2A-H2B exchange between nucleosomes. (A) The fractions of H2A-H2B exchanged nucleosomes are plotted against time in unmodified nucleosomes under various conditions as are marked in the charts. The rate constants are from the fittings (red lines) with Eq. 1 (distal H2A-H2B exchanged fraction = 0.196[1-exp(-0.77*kt*)], where *k* is the rate constant and *t* is time) and the concentration of the nucleosomes (400 nM). The error bars are the standard deviations from 4 – 10 independent measurements. The same plots and fittings are shown under the same set of conditions as in A with (B) histone H3 K56 acetylated nucleosomes and (C) CpG methylated nucleosomes.

To investigate the effect of a higher monovalent salt concentration, we measured the exchange rate at 50 mM NaCl and 150 mM KCl (Fig. 3A). The rate constant is 686 ± 111 M^-1^s^-1^, corresponding to 21 ± 3 sec H2A-H2B exchange time at a 70 μM nucleosome concentration. The significant increase in the rate supports a mechanism where a stronger ionic condition facilitates transient nucleosome partial disassembly, resulting in a higher fraction of nucleosomes eligible for inter-nucleosomal DNA-histone interactions upon collision and subsequent histone exchange. This mechanistic model is depicted in figure 4.

**Figure 4.**
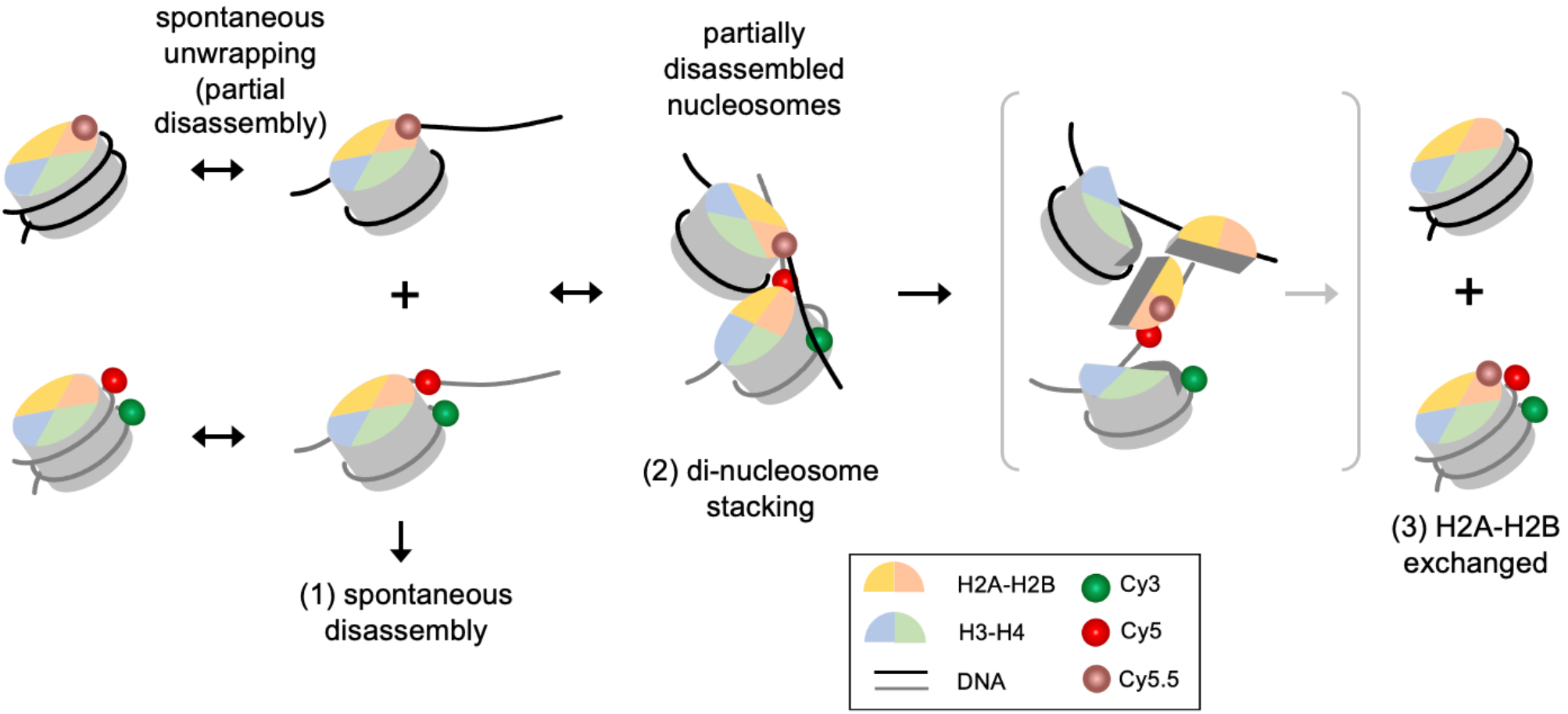
A mechanistic model for H2A-H2B exchange between nucleosomes. (1) Transient and repetitive unwrapping of nucleosomal DNA leads to spontaneous disassembly of nucleosomes at a low concentration. (2) At a high nucleosome concentration, transiently unwrapped nucleosomes may form a di-nucleosome stack via inter-nucleosomal DNA-histone interactions. (3) H2A-H2B is exchanged between two nucleosomes by the histone chaperoning function of transiently disassembled nucleosomes.

We also monitored H2A-H2B exchange at an elevated temperature of 25 °C from 4 °C. A temperature increase may destabilize DNA-histone interactions and facilitate nucleosome diffusion and nucleosome-nucleosome collisions. However, the exchange rate constant did not change much at 414 ± 64 M^-1^ s^-1^ which is within error from the value at 4 °C (Fig. 3A). This result suggests that the anticipated effects at 25 °C are not sufficient to induce a significant change in the exchange rate constant.

### Histone chaperone Nap1 facilitates H2A-H2B exchange

Histone chaperones mediate the interactions between histone and DNA, and drive them to their equilibrium, facilitating nucleosome assembly or disassembly depending on the context^38^. According to this mechanism, histone chaperones should catalyze inter-nucleosomal DNA-histone interactions as well where histones are spontaneously exchanged and redistributed among nucleosomes. We monitored H2A-H2B exchange in the presence of Nap1 at 400 nM at 4 °C. The exchange rate constant is 683 ± 94 M^-1^s^-1^ (Fig. 3A) which is significantly higher than the rate without Nap1 at 356 ± 64 M^-1^s^-1^. As the action of Nap1 requires partially disrupted DNA-(H2A-H2B) contacts, this result supports the existence of the partially disassembled nucleosomes for spontaneous histone exchange (Fig. 4). This result also supports that histone exchange may be facilitated by histone chaperone *in vivo* as various histone chaperones are abundant in the nucleus.

### H3K56 acetylation facilitates H2A-H2B exchange

Histone acetylation is typically associated with elevated gene activities^7,17,30,39,40^. In particular, acetylation at H3 K56 (H3K56ac) has been coupled to active transcription and increased termini breathing dynamics of the nucleosome^7,28,39^. These results lead to a hypothesis where H3K56ac increases the fraction of transiently disassembled nucleosomes eligible for histone exchange upon collision, thereby facilitating histone exchange. We tested this hypothesis with nucleosomes with H3K56 acetylation under various conditions (Fig. 3B).

Under all the conditions tested, the rate of H2A-H2B exchange is very high with H3K56ac, reaching the measurement limit with large errors. The rates are (1.00 ± 0.20) × 10^3^ M^-1^s^-1^, 916 ± 139 M^-1^s^-1^, (1.04± 0.18) × 10^3^ M^-1^s^-1^, and (1.38 ± 0.17) × 10^3^ M^-1^s^-1^ at a low salt concentration (10 mM NaCl), at a high salt concentration (50 mM NaCl + 150 mM KCl), at an elevated temperature (25 °C), and in the presence of Nap1, respectively (Fig. 3B). The only condition that did not show a significantly higher rate is at the high salt concentration likely because the rate is already high without H3K56ac (916 ± 139 M^-1^s^-1^ and 686 ± 111 M^-1^s^-1^ for acetylated and unacetylated nucleosomes, respectively) (Fig. 3). Under all the conditions tested, the rates reached nearly the limit of the measurements as the change is almost complete at the third time point (i.e. 2 h time point with 1 h min separation between two time points). As a result, no significant difference could be detected among the reactions under various conditions. Regardless, it is clear that H3K56ac promotes H2A-H2B exchange likely via the same mechanism as in the above cases. Under this mechanism, H3K56ac weakens the DNA-histone interactions near the termini by removing a positive charge on the histone surface, thus facilitating spontaneous partial disassembly of the nucleosome.

### CpG methylation inhibits H2A-H2B exchange

CpG methylation has been coupled to gene silencing^41^. Previous studies reported a more compact structure of CpG methylated nucleosomes toward the nucleosome termini^21,22^. Increased DNA compaction around the histone core would lower the fraction of kinetically unstable nucleosomes, which will decrease the efficiency of spontaneous H2A-H2B exchange. We monitored histone exchange with CpG methylated nucleosomes under the same salt and temperature conditions as in those for the unmodified nucleosomes (Fig. 3C).

The results indicate that CpG methylation under a physiological salt condition at 50 mM NaCl and 150 mM KCl inhibits histone exchange considerably at 397 ± 51 M^-1^s^-1^ as compared to the rate without methylation at 686 ± 111 M^-1^s^-1^, which is in line with previous studies suggesting tighter nucleosome wrapping^21^. The difference is insignificant at a low salt concentration of 10 mM NaCl (277 ± 42 M^-1^s^-1^ and 356 ± 64 M^-1^s^-1^ for the methylated and non-methylated cases) likely due to the already low rate of exchange without methylation. No difference is detected at an elevated temperature at 25 °C (400 ± 53 M^-1^s^-1^ and 414 ± 64 M^-1^s^-1^ for the methylated and non-methylated cases), suggesting that the anticipated changes are not large enough to affect the exchange rate. Despite CpG methylation suppressing histone exchange, Nap1 mediates histone exchange among CpG methylated nucleosomes as efficiently as among unmodified ones (626 ± 72 M^-1^s^-1^ and 683 ± 94 M^-1^s^-1^ for the methylated and unmethylated cases). This result suggests that the action of histone chaperone in facilitating histone exchange in chromatin is not inhibited by CpG methylation.

## DISCUSSIONS

The nucleosome is a dynamic structure that allows access to the DNA wrapped around the histone core while keeping the chromatin structure intact and suppressing faulty genome transactions. It has been reported that nucleosomes are stable under physiological ionic conditions only at above certain threshold concentration *in vitro*^3,6^. At a lower concentration, nucleosomes spontaneously disassemble within a few hours^6,42^. An unanswered question is how these fragile nucleosomes are stable at a high concentration such as in chromatin, which allows for tight packaging of genes while they are still accessible when needed. We set out to test if nucleosomes can efficiently act as their own histone chaperone and protect themselves from disassembly, while the DNA-histone contacts are repeatedly, transiently, and partially disrupted all the time.

Histone chaperones can mediate assembly or disassembly of the nucleosome depending on the context, and eventually drive a DNA-histone mixture to their thermodynamic equilibrium^10^. Histone chaperones typically contain an acidic region, enabling them to compete against DNA for histone binding^12,13^. According to the mechanism, it is reasonable to hypothesize that naked DNA regions and transiently unwrapped DNA regions from the nucleosome should also act as histone chaperone. Such a chaperone function will result in the protection of the nucleosome from disassembly at the cost of freely exchanged histones. We investigated whether nucleosomes spontaneously exchange histone H2A-H2B and if so, how fast histone exchange occurs.

Our results indicate that nucleosomes spontaneously exchange histone H2A-H2B. The rate is 356 M^-1^s^-1^ at 10 mM NaCl and is elevated to 686 M^-1^s^-1^ at 50 mM NaCl and 150 mM KCl. This increase in the rate supports that histone exchange takes place via partially unwrapped nucleosomal DNA serving as histone chaperone for another nucleosome with partially disrupted DNA-(H2A-H2B) contacts. A stronger ionic condition will increase the fraction of partially unwrapped nucleosomes and, subsequently, the chance of histone exchange upon their collisions. To support the di-nucleosomal state of this exchange mechanism (Fig. 4), stacking between a partially disassembled nucleosome and an intact nucleosome has been reported^9^. Our results with Nap1-mediated histone exchange also support this mechanism where Nap1 stabilizes partially disassembled nucleosomal states via its interactions with histone, thereby facilitating histone exchange^43^.

Our results indicate that CpG methylation and histone H3K56ac meaningfully change the rate of histone exchange under a physiological ionic condition. CpG methylation is strongly tied to gene silencing^44,45^. According to our results, CpG methylation may contribute to keeping quiet regions of the genome intact by suppressing transient partial disassembly of the nucleosome and subsequent histone exchange. The result is in line with a previous report supporting tighter DNA wrapping around the nucleosome by CpG methylation^21,31^. As for the effect of H3K56ac which is coupled to active transcription, it promotes H2A-H2B exchange as is supported by our previous reports on facilitated nucleosome termini opening motions ^22,29^. All of these results further support that histone exchange takes place via a di-nucleosomal state where partially disassembled nucleosomes form inter-nucleosomal DNA-histone interactions (Fig. 4).

Our results suggest that a histone H2A-H2B dimer is exchanged between two nucleosomes within 21 ± 3 sec under a physiological ionic condition at 70 μM nucleosome concentration which is an approximate level of nucleosome concentration in a human nucleus. This is a fairly high rate by which histones diffuse among nucleosomes in a compact chromatin context and locally concentrated histone variants and modifications of H2A-H2B will be diluted away. This spontaneous histone exchange might be a mechanism to remove gene-specific modifications of H2A-H2B after they serve their purpose. In this mechanism, H2A-H2B modifications and variants will be diffused over a large area of chromatin and eventually reverted by enzymes with no gene specificity.

## DATA AVAILABILITY

All data are contained within the manuscript.

## ACKNOWLEDGMENTS

This research was funded by NIH grants (R01 GM123164 and R01 GM130793) to T.L.

## Notes

### Competing Interest Statement

The authors have declared no competing interest.

