## Supporting Information for "Spontaneous Histone Exchange Between Nucleosomes"

**Table S1.** The sequences of DNA oligonucleotides used to construct the nucleosomal DNA. For Cy3/Cy5 labeling along the DNA, the color-marked modified nucleotides in red and green, respectively for Cy3 and Cy5, were utilized. iAmMC6T is a Thymine analog with an amine terminated C6 linker is attached.

| | | |
|---|---|---|
| F1 | 34 | /5Biosg/GCAG ATCGAGAATC CCGGTGCCGA GGCCGCTCAA |
| F2 | 77 | /5Phos/ TTGG /iAmMC6T/CGTAGACAG CTCTAGCACC GCTTAAACGC ACGTACGCGC TGTCCCCCGC GTTTTAACCG CCAAGGGGAT TAC |
| F3 | 40 | /5Phos/ TCC C/iAmMC6T/AGTCTCCA GGCACGTGTC AGATATATAC ATCCGAT |
| R1 | 69 | ATCGGATGTA TATATCTGAC ACGTGCCTGG AGACTAGGGA GTAATCCCCT TGGCGGTTAA AACGCGGGG |
| R2 | 78 | /5Phos/G ACAGCGCGTA CGTGCGTTTA AGCGGTGCTA GAGCTGTCTA CGACCAATTG AGCGGCCTCG GCACCGGGAT TCTCGAT |

**Table S2.** The sequences of DNA oligonucleotides used to construct the DNA mimetics for the two different H2A-H2B exchange cases. The labeling positions of Cy3, Cy5, and Cy5.5 are denoted in green, red, and brown, respectively. The mark iAmMC6T is for a Thymine analog with an amine terminated C6 linker.

| Distal Case | |
|---|---|
| S1 | /5 biosg/ ACGCGAGC/iAmMC6T/CAGGAGCA |
| S2 | /5Cy5/TGCTCCTGAGCTCGCGT/3Cy55Sp/ |
| Proximal Case | |
| S3 | /5 biosg/ ACGCGAGC/iAmMC6T/CAGGAGCA |
| S4 | /5Cy55/TGCTCCTGAGCTCGCGT/3Cy3Sp/ |

**Table S3.** The fitting results from the charts shown in figure 3. The apparent kinetic rate constants are in $h^{-1}$ and not corrected for the nucleosome concentration.

| | **4 °C, 10 mM NaCl ($h^{-1}$)** | **4 °C, 50 mM NaCl + 150 mM KCl ($h^{-1}$)** | **25° C, 10 mM NaCl ($h^{-1}$)** | **+ Nap1, 4°C ($h^{-1}$)** |
|---|---|---|---|---|
| **Unmodified** | 0.513 ± 0.092 | 0.989 ± 0.158 | 0.595 ± 0.090 | 0.982 ± 0.137 |
| **CpG methylated** | 0.396 ± 0.059 | 0.570 ± 0.072 | 0.574 ± 0.077 | 0.901 ± 0.104 |
| **H3K56ac** | 1.44 ± 0.28 | 1.32 ± 0.20 | 1.50 ± 0.26 | 1.97 ± 0.24 |

**Figure S1.** Native PAGE analyses for nucleosomal DNA and nucleosomes. (A) The left two lanes ($L_{DNA}$ and $U_{DNA}$) show Cy3/Cy5 labeled and unlabeled nucleosomal DNA, respectively, and the right two lanes ($L_{Ac}$ and $U_{Ac}$) show H3 K56 acetylated nucleosomes with labeled and unlabeled DNA, respectively. (B) Lanes $L_{un}$ and $U_{un}$ show unmodified nucleosomes with labeled and unlabeled DNA. Lanes $L_{MT}$ and $U_{MT}$ show CpG methylated nucleosomes with labeled and unlabeled DNA. Note that the nucleosomes with unlabeled DNA have their H2B labeled with Cy5.5.

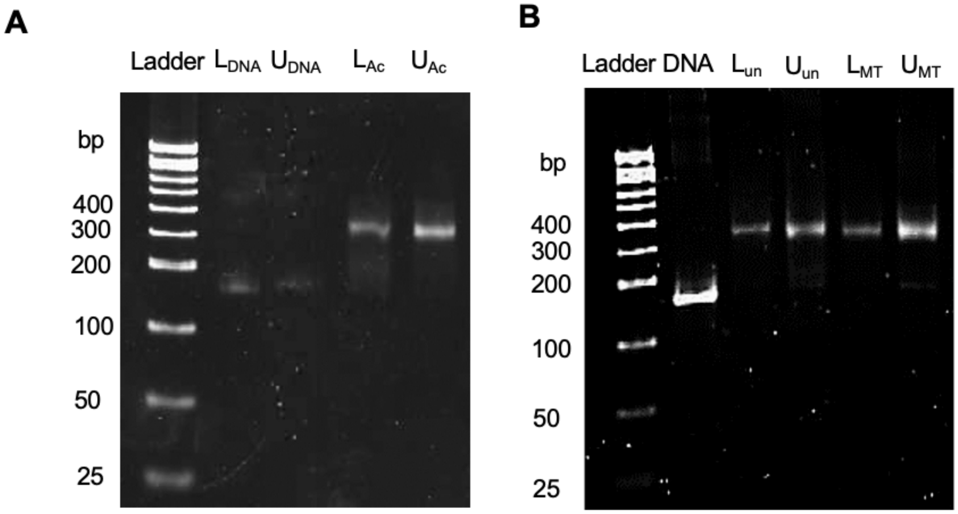

**Figure S2.** Control experiments to confirm fluorescence intensity signatures from H2A-H2B exchanged nucleosomes. (A) The structure of a nucleosome core particle with the labeling positions marked (left). The estimated distances between the fluorophores are shown (right). (B) The structures of 15 bp biotinylated dsDNA labeled with fluorophores at the marked positions that can mimic the H2A-H2B exchanged nucleosomes according to the distances estimated in A. (C) Intensity traces from fluorophores labeled at DNA mimetics are shown in B. In case of the proximal H2A-H2B exchange mimetic (left), a high level of Cy5.5 and a low level of Cy3 and Cy5 signals were observed, followed by photobleaching. In case of the distal H2A-H2B exchange mimic (right), a low level of Cy3 and a mid-high level of Cy5 and Cy5.5 signals were observed.

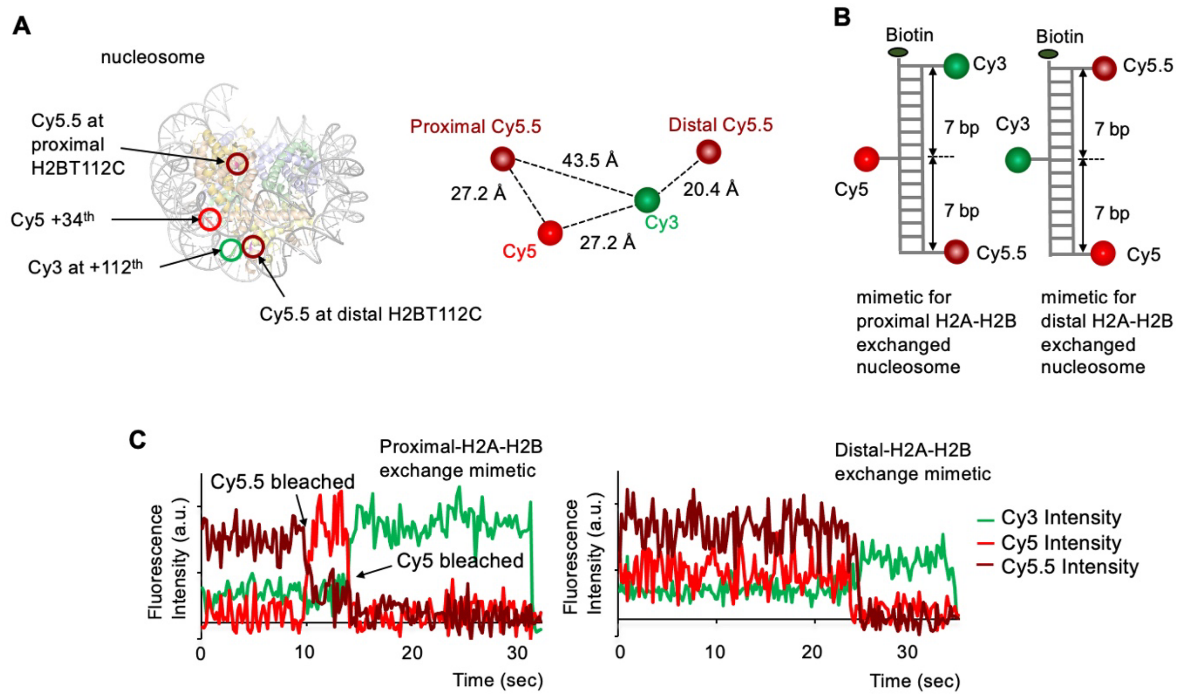

**Figure S3.** Nucleosomes with different labeling possibilities during the histone exchange reaction. (A) A total of eight different nucleosome species with a different fluorophore labeling scheme can exist in the reaction mix. (B) The full list of possible combinations of histone exchange between the eight species shown in A.

**A**

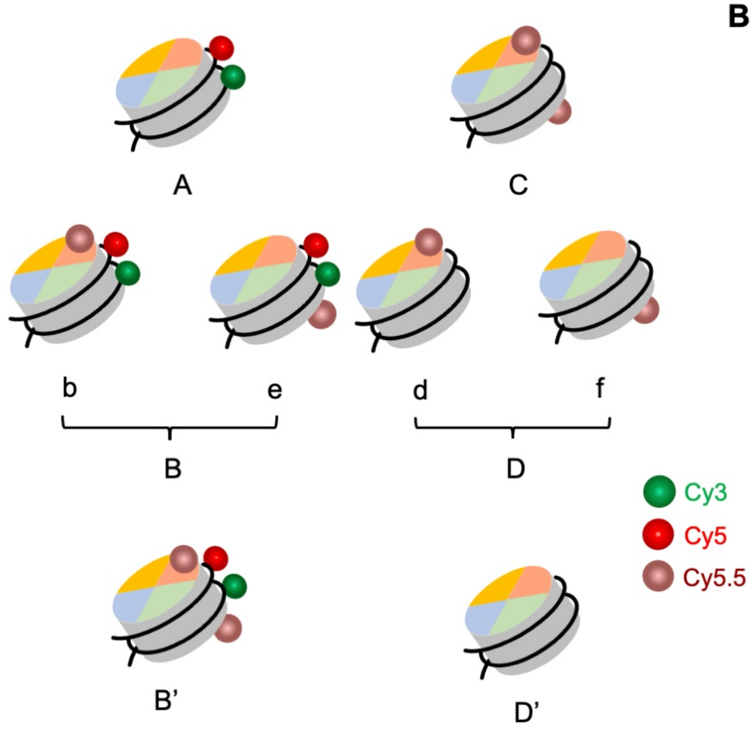


**B**

| | | |
|---|---|---|
| A+C | → | B+D |
| A+D | → | B+D’ |
| | → | A+D |
| A+D’ | → | A+D’ |
| B+C | → | B+C |
| | → | B’+D |
| B+D | → | 2(B+D) |
| | → | B’+D’ |
| | → | A+C |
| B+D’ | → | B+D’ |
| | → | A+D |
| A+B | → | 2(A+B) |
| C+D | → | 2(C+D) |
| C+D’ | → | D+D |
| D+D’ | → | D+D’ |
| A+A | → | A+A |
| B+B | → | B+B |
| | → | A+B’ |
| C+C | → | C+C |
| D+D | → | D+D |
| | → | D’+C |
| D’+D’ | → | D’+ D’ |

**Figure S4.** Histograms of $FRET_{Cy5}$ and $FRET_{Cy5.5}$ at 0 and $18^{th}$ hour time points from the nucleosomes showing zero intensities after photobleaching (single nucleosomes for accurate background correction), longer photobleaching lifetimes (> 10 sec), and a decent signal-to-noise ratio (> 4) for (A) unmodified, (B) histone H3K56 acetylated (0 and $8^{th}$ hour), and (C) CpG methylated nucleosomes. It is evident in all cases that both non-zero $FRETCy_{5.5}$ and lower $FRET_{Cy5}$ grow as time passes (see red-circled areas), indicating that H2A-H2B-Cy5.5 is being incorporated into the nucleosomes labeled with a Cy3-Cy5 pair. Note that the histogram counts cannot be used as a measure of the percent exchange as Cy5.5 photobleaches much faster than Cy5. It is typical according to our experience that fluorophores labeled on protein photobleach much faster than those labeled on nucleic acids.

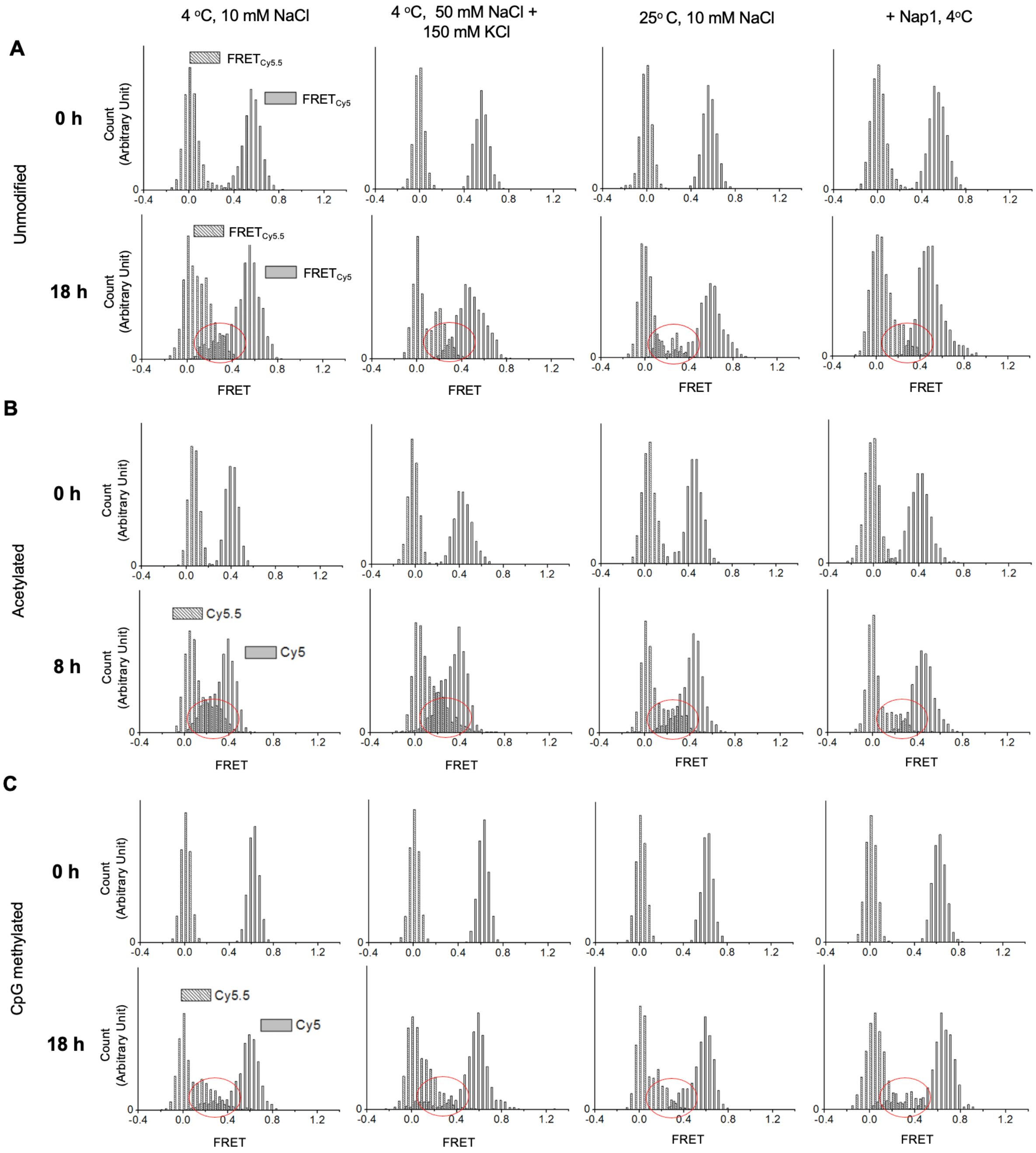
